# Metaproteomics revealing cyanide bioremediators in effluents from artisanal cassava processing in the Brazilian Amazon

**DOI:** 10.1101/2025.05.20.655124

**Authors:** Alana Coêlho Maciel, Sayure Mariana Raad Nahon, Isa Rebecca Chagas da Costa, Rafael Borges da Silva Valadares, Hellen Kempfer Philippsen, Celina Eugenio Bahule, Beni Jequicene Mussengue Chaúque, Kelton Henrique Alves Guimarães, Nayara Macêdo Peixoto Araújo, Alessandra Santos Lopes

## Abstract

Cassava is widely used and processed for food in several countries. During artisanal processing for flour production, liquid waste containing cyanide may be disposed of inappropriately, contaminating soils, rivers and streams. Various microorganisms must proliferate and carry out fermentation from this waste. The study was carried out on a family farm in the municipality of Bragança, in the state of Pará, Brazil. Samples were selected from the effluents generated in four stages (fermentation tank, post-fermentation washing water tank, washing/pressing water tank (*manipueira*) and effluent lagoon). The aim of this study is therefore to track down microorganisms and their enzymes, through metaproteomics, at all stages of cassava processing up to the elimination of wastewater discharges, using the metaproteomics technique in order to track down potential cyanide bioremediators adapted to local environmental conditions. The samples were extracted using the Phenol-SDS protocol, washed, digested and submitted to a high-resolution mass spectrometer coupled to liquid chromatography and subsequently analyzed using sensitive software. Microorganisms with potential cyanide bioremediation potential were identified (*Bacillus subitilis, Pseudomonas putida* and *Levilactobacillus brevis*), as well as nitrogen fixers involved in bioremediation. The identified peptides analyzed showed proteins involved in: DNA repair; glycosylation; phosphorylation; CAZY proteins, among others. These findings provide a foundation for clean biotechnological applications aimed at treating cyanide-contaminated effluents in artisanal cassava processing in the Amazonian territory.

**Importance:** This study is important due to the future perspectives for bioremediating cyanide, identifying active enzymes and metabolic pathways, discovering and optimizing consortia of native key microorganisms, monitoring treatment efficiency in real time, having sustainable environmental and human health alternatives, valuing traditional knowledge and regional development in the Amazonian context.

**Key Points:** Prospecting for bioremediators generated from cassava effluents;

Metaproteomics clarified the microbial dynamics present in cassava effluents;

Metaproteomics identifying active enzymes and metabolic functions in biotechnology.

## Introduction

Cyanide, a highly toxic nitrile, poses significant health risks to both humans and animals. It can be produced through natural and anthropogenic processes and is widely used in industries such as jewelry, mining, electroplating, plastics, dyes, paints, pharmaceuticals, and coal coke production. Residues from these processes are often discarded into the environment (Sharma, Akhter, and Chatterjee, 2019). Similarly, in cassava processing, cyanide-containing residues are generated through the degradation of cyanoglycosides and cyanolipids found in the plants, particularly in rural Amazonian communities where effluent management remains informal or absent. In the Legal Amazon region of Pará, traditional communities cultivate and process cassava roots (*Manihot esculenta* Crantz) to produce products like flour, *tapioca, tarubá*, and *tucupi*. During processing, cyanide residues are frequently released into effluents and soils, posing a threat to environmental, animal, and human health (Maciel et al., 2023). Cyanide contamination can lead to the formation of various compounds and complexes, some more toxic than others, including metal-cyanide complexes, thiocyanates, and nitriles. These reactions are influenced by factors such as pH, temperature, oxygen availability, ammonia and metal ions (Maciel et al., 2023; Malmir et al., 2022).

Microorganisms, such as fungi and bacteria, can both produce cyanide during metabolic activities through the cyanogenesis process, and can also degrade it through various biochemical pathways. These include reductive, oxidative, hydrolytic, and substitution/transfer reactions (Mekuto et al., 2016).

Bioremediation through microbial degradation of cyanide has proven successful in several studies and is recommended for environments impacted by cyanide, including effluents, rivers, ponds, cattle drinking troughs, and agricultural soils (Razanamahandry et al., 2017). For efficient bioremediation, microorganisms require easily metabolizable substrates like glucose and sucrose. Additionally, enzymatic degradation strategies using microbial and plant enzymes have been adopted. These enzymes include cyanidase, cyanide hydratase, formamidase, nitrilase, nitrile hydratase, cyanide dioxygenase, cyanide monooxygenase, cyanide cyanase, and nitrogenase, all of which have been extensively studied (Seteno, Ntwampe, Bruno, 2014). Identified microorganisms likely to be effective in bioremediating cyanides include *Pseudomonas* sp., *Acinetobacter* sp., *Aspergillus* sp., *Pseudomonas pseudoalcaligenes, Azotobacter chroococcum* NCIMB 8003, *Azotobacter vinelandii, Alcaligenes xylosoxidans* subsp. *denitrificans* DF3, *Bacillus pumilus* C1, *Pseudomonas fluorescens* NCIMB 11764, and *Pseudomonas stutzeri* AK61 (Sharma, Akhter, Chatterjee, 2019; Cabello et al., 2018).

Advances in omics technologies, such as metaproteomics, have been instrumental in identifying microorganisms and enzymes involved in cyanide bioremediation and environmental recovery. For example, metaproteomics has provided detailed insights into the microbial profiles and enzymes involved in processes like corn fermentation, facilitating the selection of optimal process starters (Bahule et al., 2023; Yang, Fan, Xu, 2020). Another study using metaproteomics analyzed effluent composition, microbiota, and proteins associated with stress responses, cellular structures, and metabolic functions, offering new perspectives on agriculture, environmental quality recovery, biotechnological process control, and sustainable measures (Armengaud, 2023).

Omics-based research on cassava has highlighted microorganisms with high metabolic capacities that transform or degrade chemical contaminants into non-toxic or less hazardous products (Sharma et al., 2022). Additionally, it has identified beneficial microorganisms in the soil associated with cassava roots, which enhance plant productivity and resistance (Benko-Iseppon et al., 2021; Ding et al., 2020).

This study aims to characterize the metaproteomic profile of microbial communities across cassava processing stages and identify key taxa and enzymes involved in cyanide degradation, with potential application in bioremediation strategies.

## Materials and methods

### Sampling of cassava processing effluents

Liquid waste samples from cassava processing were collected in a rural area of Bragança (1°02’06.6”S; 46°49’42.8”W), Pará, Brazil, in March 2023. Collections took place at four key stages of the artisanal cassava processing: (1) Fermentation tank (FT) – where whole tubers are submerged for 3 days; (2) Post-fermentation wash water tank (WWT) – where peeled tubers are soaked for 1 day; (3) Washing/pressing water tank (PW) (*manipueira*) (WP); (4) Effluent lagoon (EL) – the final destination for liquid waste generated during processing. Samples were stored in sterile 1000 ml polyethylene bottles without preservatives and transported to the laboratory within 6 hours of collection. They were kept at 2 to 5 °C until metaproteomic analysis was performed.

### 2.2 Determination of total and free hydrocyanic acid

The quantification of total and free hydrocyanic acid (HCN) was performed following the method described by Modesto et al. (2019), with slight adaptations. Briefly, to determine the total HCN, 0.4 ml of buffer solution (pH 7.0) was added to a test tube, followed by 0.1 ml of each sample (in triplicate) from the fermentation tank, washing tank, pressing water, and effluent lagoon. Then, 0.1 ml of β-glucosidase (4 U, commercial lyophilized almond enzyme, Sigma-Aldrich) was added. The test tubes were heated at 30 °C in a water bath for 15 minutes to hydrolyze the cyanogenic compounds. After heating, 0.6 ml of 0.2 M NaOH was added to fix the HCN in the solution and prevent its volatilization, and the tubes were left to stand for 5 minutes. Subsequently, 2.8 ml of buffer solution (pH 6.0) and 0.1 ml of chloramine T were added, and the tubes were placed in an ice bath (≈ 5°C) for 5 minutes. After this time, 0.6 ml of color reagent (a solution of 1,3-dimethyl barbituric acid and isonicotinic acid) was added, and the solution was vortexed. The absorbance was then measured using a spectrophotometer (Thermo Scientific, Evolution 60, Massachusetts, USA) at 605 nm after incubation at room temperature for 10 minutes.

For the determination of free cyanide (CN-), the same procedure was followed, but without the addition of β-glucosidase. Instead, 3.4 ml of buffer solution (pH 6.0) was added, followed by 0.6 ml of each sample in triplicate (from the fermentation tank, washing tank, pressing water, and effluent lagoon), and 0.1 ml of chloramine T. The tubes were then placed in an ice bath (≈ 5°C) for 5 minutes. Afterward, 0.6 ml of color reagent was added, and the absorbance was measured at 605 nm after 10 minutes at room temperature. An analytical curve for quantification was constructed using concentrations ranging from 0.5 to 10.0 μg of HCN/ml (r^2^ = 0.9941) for both total and free cyanide.

### Metaproteomics

### Protein extraction

Protein extraction was carried out following the protocol described by Wang et al. (2006), with modifications by Trindade et al. (2021). Briefly, approximately 40 ml of liquid effluent from each sample was transferred to 50 ml Falcon tubes and centrifuged at 9,000 rpm for 6 minutes at 4°C. The supernatant was discarded. Then, 3.0 ml of extraction buffer [0.85 M sucrose, 0.1 M Tris-Hydrochloride (Tris-HCl, pH 8.0), 2% (w/v) sodium dodecyl sulfate (SDS), 1 mM phenylmethylsulfonyl fluoride (PMSF), and 2% (w/v) polyvinylpolypyrrolidone (PVPP)] was added, along with 210 μl of dithiothreitol (DTT) and 3 μl of Protease Inhibitor Cocktail Powder (Merck, St. Louis, MO, USA). The samples were incubated at room temperature for 10 minutes and sonicated five times for 30 seconds at 70 watts.

Each sample was then divided into 1 ml aliquots in two microtubes, and 700 μl of phenol was added to each. The samples were centrifuged at 14,000 rpm for 7 minutes at 4°C. The phenolic phase from each aliquot was collected and combined into new microtubes, followed by another centrifugation at 14,000 rpm for 7 minutes at 4°C to remove any residual SDS or aqueous phase. The phenolic phase was collected, and proteins were precipitated by incubating the samples with 550 μl of 0.1 M ammonium acetate (prepared in absolute methanol at −20°C) at −80°C overnight.

### Sample Washing

The samples were centrifuged at 14,000 rpm for 7 minutes at 4°C, and the supernatant was discarded. The protein pellets were washed twice with 1.5 ml of ice-cold 80% (v/v) acetone and centrifuged at 14,000 rpm for 4 minutes at 4°C each time. After discarding the acetone, 1.5 ml of ice-cold 70% (v/v) ethanol was added, and the samples were centrifuged again at 14,000 rpm for 4 minutes at 4°C. Finally, 100 μl of RapiGest™ Surfactant (0.1%, Waters) was added, and the samples were stored at -80°C until protein digestion. Protein quantification was performed using the Qubit 3.0 Fluorometer (Invitrogen).

### Protein digestion

The proteins were reduced with 5 mM Dithiothreitol (DTT) and incubated for 25 minutes at 56 °C. Alkylation was performed with 14 mM Iodoacetamide (IAA) for 30 minutes at room temperature, protected from light. The excess IAA was reduced by adding 5 mM DTT and incubating for 15 minutes at room temperature. Next, 1 mM calcium chloride (CaCl_2_) was added, and trypsin (50 ng/µL) treatment was carried out for 20 hours at 37 °C with shaking at 200 rpm. The enzymatic reaction was stopped by adding trifluoroacetic acid to a final concentration of 0.4%. The samples were incubated at 37 °C for 90 minutes, then centrifuged at 14,000 rpm for 10 minutes at 6 °C. The supernatants were transferred to vials, and the pH was adjusted to 10 with 1 M ammonium hydroxide to facilitate effective capture on the first-dimension column of Ultra-Performance Liquid Chromatography (HPLC). The HPLC system was directly coupled to the ESI-Q-ToF Synapt G2S mass spectrometer (Waters), operating in positive mode with continuous fragmentation (MSE) and a collision energy oscillating between 5 and 40 eV.

### Protein identification and data analysis

Protein identification and quantification were performed using Progenesis QI for Proteomics v4.2 (Nonlinear Dynamics, Waters). The search was conducted against a customized database combining Manihot esculenta protein sequences with bacterial and fungal protein sequences retrieved from the NCBI non-redundant (nr) database. The database was constructed to reflect the complexity of plant–microbe interactions present in cassava-processing environments. Searches were performed with a precursor mass tolerance of 10 ppm and fragment mass tolerance of 20 ppm. Carbamidomethylation of cysteine was set as a fixed modification, and oxidation of methionine as a variable modification. Only peptides with a false discovery rate (FDR) ≤ 1% were considered for downstream analysis.

## Results and discussion

Our results (Figure 1) show a diverse range of microorganisms present at different stages of M. esculenta processing. The majority of species (170) were detected in the fermentation tank, with particular emphasis on the genera *Mucilaginibacter* spp. (a group of endophytic bacteria commonly found in plant roots and soil; Kim et al., 2012; Zhou et al., 2019) and *Mycena* spp. (small, primarily saprophytic mushrooms associated with the roots of living or decaying plants; Christoffer et al., 2023).

**Figure 1.**
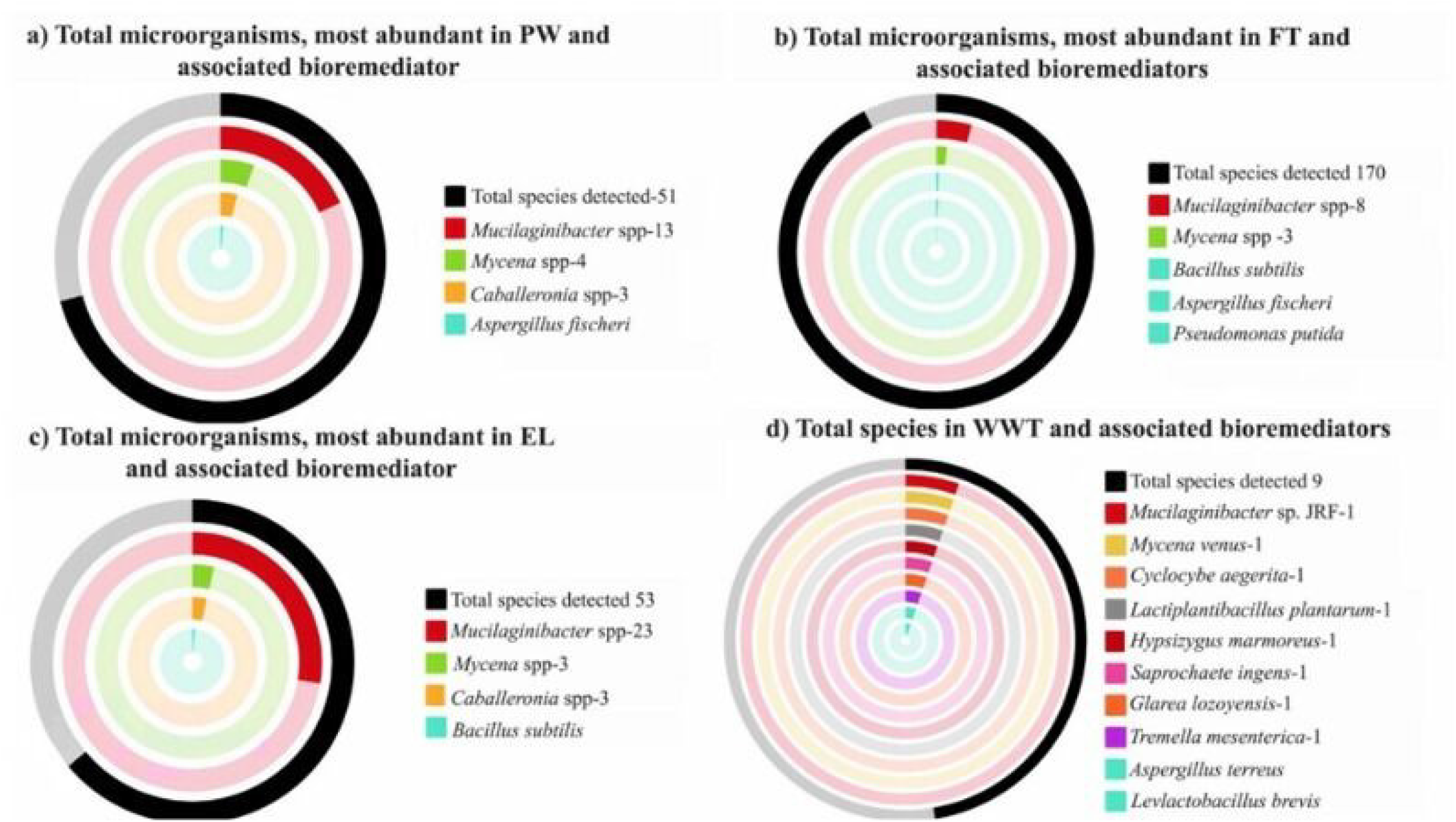
Diversity of microorganisms at different stages of cassava processing: Fermentation tank (FT), Post-fermentation washing water tank (WWT), Washing/pressing water tank (PW), and Effluent lagoon (EL))

**<Figure 1, near here.>**

As shown in Figure 1, Mucilaginibacter and Mycena spp. were consistently abundant across all stages, although not previously described as cyanide degraders. The genera *Mucilaginibacter spp*. and *Mycena spp*. were present throughout all stages of cassava processing, including fermentation, washing, pressing, and disposal. It is important to note that, despite their abundance, these genera have not been previously reported as cyanide bioremediators.

Among the various genera identified, five species have been established as having cyanide bioremediation capabilities: *Aspergillus fischeri, Aspergillus terreus, Bacillus subtilis, Levilactobacillus brevis*, and *Pseudomonas putida*. The microorganisms identified in cassava processing and associated with cyanide bioremediation activity are listed in Table 1.

**Table 1.** Cyanide bioremediating microorganisms and the total and free cyanide concentrations at different stages of cassava processing.

| Sampling site | Total HCN (mg L<sup>-1</sup>) | Free HCN (mg L<sup>-1</sup>) | Microorganisms found |
| --- | --- | --- | --- |
| Fermentation tank (FT) | 1.66±0.04 | 0.20±0.02 | *Bacillus subtilis*; *Aspergillus fischeri*; *Pseudomonas putida* |
| Post-fermentation Washing water tank (WWT) | 7.21±0.48 | 0.14±0.00 | *Aspergillus terreus*; *Levilactobacillus brevis* |
| Washing/pressing water tank (PW) | 5.58±0.33 | 0.79±0.04 | *Aspergillus fischeri* |
| Effluent Lagoon (EL) | 1.39±0.37 | 0.07±0.00 | *Bacillus subtilis* |

The data (Table 1) show that *B. subtilis* was detected during both cassava fermentation and in the effluent lagoon, suggesting that it multiplies during fermentation and withstands the conditions of cassava processing. *A. fischeri* was identified in both the fermentation tank and during pressing. *P. putida* occurred only during fermentation, while *A. terreus* and *L. brevis* were found in the post-fermentation wash water. The microorganisms identified by metaproteomics (*A. fischeri, A. terreus, B. subtilis, L. brevis*, and *P. putida*) are known cyanide degraders, as reported in environmental or in vitro studies (Meyers et al., 1993; Potot, 2010). The metaproteomic data confirmed not only the presence, but the metabolic activity of these species via expressed peptides associated with known cyanide degradation pathways.

*B. subtilis* is a well-studied model organism within Gram-positive bacteria, widely researched for its biochemical, genetic, and molecular properties (Manabe et al., 2011; Vojcic et al., 2012). It has been primarily used in industrial enzyme and metabolite production, as well as biological control (Dischinger et al., 2009; Yanjie et al., 2022). In the context of cyanide degradation, *B. subtilis* and *B. pumilus* degraded 100% of cyanide at 27°C and 65 rpm in in vitro cultures. Other *Bacillus* species, such as *B. licheniformis*, have also been reported to degrade cyanide, achieving 99.5% degradation at 35°C and 190 rpm (Meyers et al., 1993). Other studies have shown that bacterial consortia of *B. subtilis* and *P. stutzeri* can degrade cyanide in soils contaminated by cyanide from cassava processing (Nwokoro, Dibua 2014).

*P. putida*, another bacterial species known to degrade cyanide in cassava waste, breaks down cyanide into ammonia. This species is a nitrogen fixer, consumes by-products from biodiesel waste, inhibits pathogenic bacteria, and is used in biotechnology for producing gallic acid and expressing the enzyme nitrile hydratase (Dias et al., 2022; Weimer et al., 2020; Verhoef et al., 2014; Watanabe et al., 1998; Park et al., 2017). Additionally, *P. putida* has been identified alongside *Patotea agglomerans* as a phosphorus-solubilizing bacterium in latosol (Junior Barbosa, Gentil, 2023). Given its ability to degrade cyanide and its presence in cassava processing effluents, *P. putida* shows promise for intentional bioremediation.

*L. brevis*, another bacterium found in the fermentation tank, is known to degrade cyanide. It degraded approximately 37% of cyanide during fermentation of two cassava varieties (*kibandameno* and *tajirika*) and enhanced the plant’s resistance to environmental stress (Okoth et al., 2022).

*Aspergillus terreus*, isolated from low pH soil, thrives across a range of pH (4–7). In bioremediation studies, it removed varying amounts of copper (69%, 47%, 38%, and 24% for 100, 200, 300, and 500 mg/L concentrations, respectively) and effectively bioremediated lead (Pb), cadmium (Cd), chromium (Cr), and nickel (Ni) in wastewater and industrial effluents (Dias et al., 2002). *A. terreus* is non-pathogenic, easy to cultivate, and highly resistant to effluent toxicity. Additionally, *A. terreus* and *A. fumigatus*, sourced from enriched effluents, bioremediated metals such as Cd (93.28%), Cr (89.41%), and Pb (97.13%) both individually and in consortia (Zango et al., 2023).

*A. fischeri*, isolated from soil samples, demonstrated significant tolerance to high copper concentrations (600 ppm), making it suitable for the bioremediation of copper-polluted environments (Amin, Nazir, 2024).

Although several enzymes associated with carbohydrate metabolism were expressed, no direct functional annotation linked them explicitly to cyanide detoxification enzymes such as cyanidases or nitrilases. Figure 2 outlines the biological processes identified based on protein functions across all processing stages. Highlighted are processes linked to energy metabolism (glycolysis, glucose metabolism), which may support microbial survival in cyanide-rich effluents.

**Figure 2.**
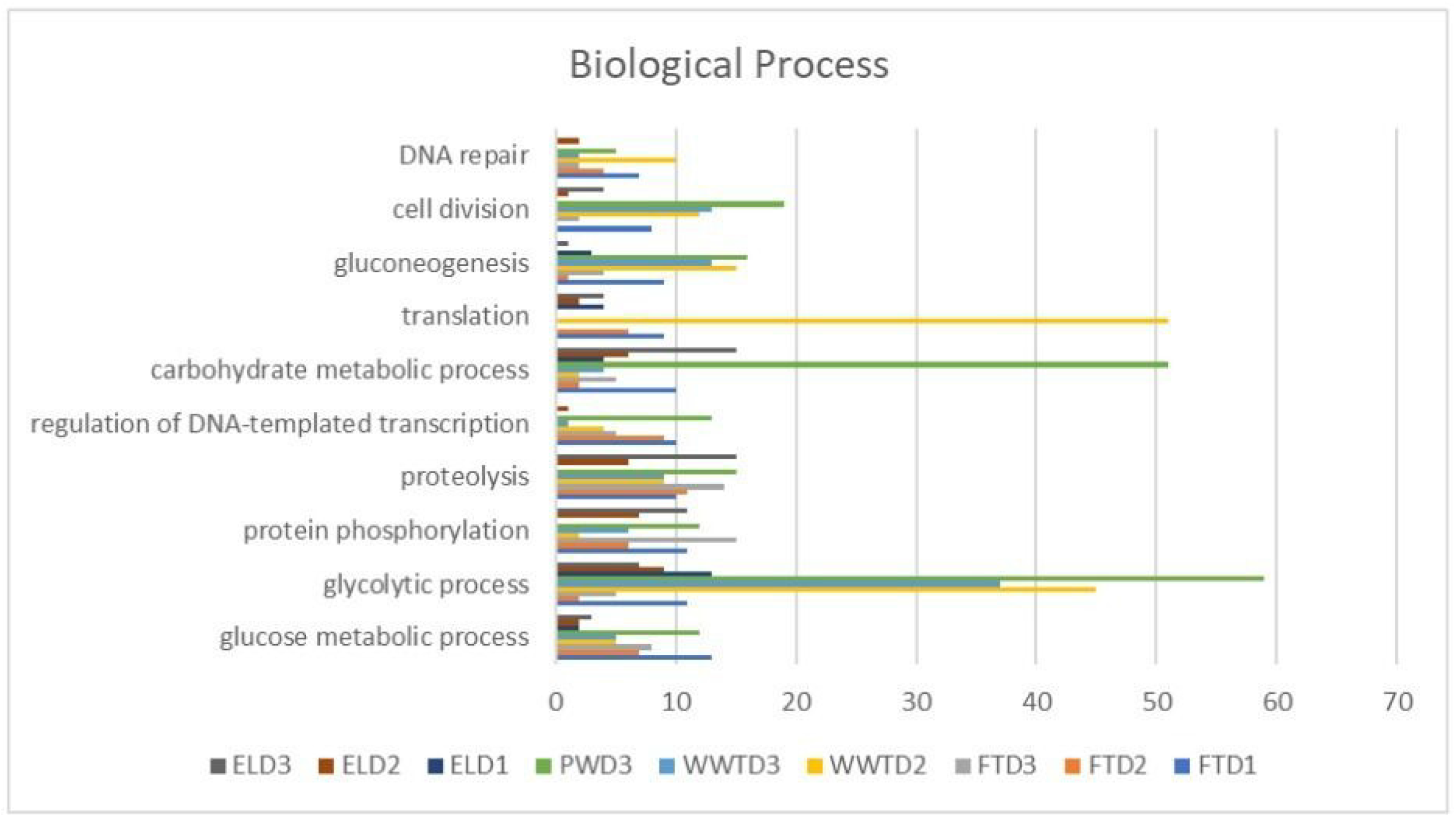
Functional category proteins: Biological processes FTD1 -Fermentation tank day 1; FTD2 -Fermentation tank day 2; FTD3 -Fermentation tank day 3; WWTD2 - Washing water tank day 2; WWTD3-Washing water tank day 3; PWD3 - Pressing water day 3; ELD1-Effluent lagoon day 1; ELD2-Effluent Lagoon day 2; ELD3-Effluent Lagoon day 3.

**<Figure 2, near here.>**

It is evident that the processes involved in the metabolism of glucose and other carbohydrates were expressed in all stages of cassava processing. This indicates that carbohydrates and glucose are available in the waste material. The presence of glucose is a fundamental condition for bioremediation, as bioremediating microorganisms require it as a carbon and energy source for their maintenance. Bioremediation is further facilitated by the presence of specific enzymes produced by various microorganisms. The persistent detection of glycolytic and carbohydrate metabolism proteins (Figure 2) across all samples suggests sustained microbial energy acquisition, a critical condition for successful bioremediation in toxic environments.

*P. agllomerans* was also identified in this study in FTD1 and FTD2. Previous studies describe it as a biofilm-forming species that creates a barrier against the penetration of toxic industrial contaminants into the soil. It also has the ability to fix N2 and can serve as a potential biofertilizer or biocontrol agent (Lorenzi et al., 2022; Toores et al., 2013). This species has been identified as a potential bioremediator of Cu (60%), Pb (96%), and Fe (96%) (Audu et al., 2020), and it also promotes plant growth (Mefteh et al., 2017).

In addition to bacteria involved in cyanide bioremediation and N2 fixation, numerous species with high carbohydrate metabolism were identified in the cassava processing environment. Some of these genera were present from the fermentation tank to the effluent lagoon, including *Burkholderia mallei, Bdellovibrio bacteriovorus, Acinetobacter baumannii, Bradyrhizobium diazoefficiens, Geobacillus stearothermophilus, Flavobacterium johnsoniae, Latilactobacillus sakei* subsp. *sakei* (strain 23K), *Neochlamydia sp*., *Sulfurovum sp*., *Enterobacter sp*. (strain 638), and *Protochlamydia amoebophila*, among others.

Although these species have not yet been reported or tested for cyanide bioremediation activity, they have demonstrated considerable resistance to environments with high cyanide concentrations. This suggests that they may play a role in bioremediation or assist with enzymes that degrade or transform cyanide into other products, which have yet to be described.

It is possible that many of the identified bacteria, which are not currently known as cyanide bioremediators, could be involved in this process, as they are capable of multiplying in environments like the effluents studied here. However, studies in controlled environments are needed to confirm this hypothesis.

Regarding the cyanide concentrations observed in the cassava processing stages, the average total cyanide in the washing water after 3 days of fermentation was high, at 7.21 mg L−1, and remained elevated at 5.58 mg L−1 in the pressing water (*manipueira*). This indicates the significant amounts of cyanide being released into the effluent lagoon. Fermentation does not appear to influence the reduction of hydrocyanic acid in any way. The results show that the levels of hydrocyanic acid, both total and free, increased as the processing stages progressed. In the effluent lagoon, the average total cyanide found was 1.39 mg L−1, a concerning value since it exceeds the maximum allowable concentration of 1.0 mg L−1 according to Conama Resolution No. 430/11 and 0.65 mg L−1 according to the EPA. For free cyanide, the permitted maximum values by Conama No. 430/11 and the EPA are 0.2 mg L−1. Although the cyanide level in the FT stage was 0.2 mg L−1 (within the legal limit), subsequent stages (WWT, PW, and EL) had levels exceeding the recommended regulations.

This highlights a clear warning for future effluents in the local environment. The data suggests that neither fermentation nor the abundance of microorganisms and enzymes found during processing influenced the reduction of hydrocyanic acid, which remained at high levels in the waste analyzed and disposed of in the effluent lagoon. The consistent presence of non-documented degraders suggests potential auxiliary roles or uncharacterized enzymatic pathways that warrant further investigation.

## Conclusions

The present study demonstrates that metaproteomics is a valuable tool for analyzing microbial communities in cassava processing, significantly reducing the time required compared to traditional culture methods. The technique provided consistent and accurate results, revealing the presence of glycolytic biological processes which is an important indicator of available carbon sources for microbial activity, a crucial condition for the bioremediation of cyanides in effluents

The findings support the hypothesis of this research, confirming the presence of a diverse range of microorganisms in cassava fermentation that are tolerant to high concentrations of hydrocyanic acid. However, their direct contribution to cyanide toxicity reduction or bioremediation remains unclear. A small subset of these microorganisms, including *A. fischeri, A. terreus, B. subtilis, L. brevis*, and *P. putida*, have already been documented for their cyanide bioremediation capabilities.

Some belong to the group of fermenters and probiotics (*B. subtilis*), are not potential food poisoning agents, and their HCN bioremediation action has already been shown in the laboratory (*P. putida*) and also in effluents (*L. brevis*).

The potential benefits for inoculating microorganisms during cassava fermentation should be considered in future bioremediation studies, as it could reduce both effluent treatment costs and fermentation time. It is possible that the activity of other microorganisms influenced the course of spontaneous fermentation and supported the survival of the observed microbial populations. Further research is needed to explore the metabolic interactions between microbial communities and their impact on cyanide reduction in effluents.

It is important to acknowledge that metaproteomics, while powerful, has limitations in the quantity and quality of protein extraction, which can influence the data generated. Therefore, caution is needed when interpreting results, especially in the context of naturally fermented foods and effluent environments, which may be influenced by various biotic and abiotic factors. Future studies should investigate the inoculation of targeted microbial consortia during fermentation to enhance both product safety and post-process effluent remediation.

## Author contributions Statement

ACM: investigation, writing - review & editing, table; SMRN; IRCC; RBSV: methods and analysis; HKP; figure, writing and discussion; CEB; writing and discussion; BJMC: discussion and graphical abstract; KHAG: figure; NMPA: validation and supervision; ASL: validation, supervision and Financing Acquisition. All authors reviewed the manuscript.

## Acknowledgments

The authors acknowledge Coordination of Higher Education Personnel Improvement (CAPES, Brazil) for providing doctoral scholarship (A.C. Maciel / Process number 88887.620123/2021-00).

## Data availability

The authors declare that the data supporting the findings of this study are available in the article and in their Supplementary Tables (ST) files.

## Consent to participate

Not applicable

## Consent to publish

Not applicable

## Ethical approval

Ethical approval was granted in accordance with the Certificate of Regularity of Access to Genetic Heritage and Associated Traditional Knowledge, registration no. AAA8C32, dated 05.06.2023, from the Genetic Heritage Management Council, Ministry of the Environment (MMA), Brazil.

## References

Amin I, Nazir R, Rather MA. (2024). Evaluation of multi-heavy metal tolerance traits of soil-borne fungi for simultaneous removal of hazardous metals. World J Microbiol Biotechnol. 40, 175. 10.1007/s11274-024-03987-z

Armengaud J. (2023). Metaproteomics to understand how microbiota function: the crystal ball predicts a promising future. Environ. Microbiol. 25, 115–125. 10.1111/1462-2920.16238

Audu KE, Adeniji SE, Obidah JS. (2020). Bioremediation of toxic metals in mining site of Zamfara metropolis using resident bacteria (Pantoea agglomerans): A optimization approach. Heliyon. 6 (8), e04704. 10.1016/j.heliyon.2020.e04704

Bahule CE, da Silva Martins LH, Chaúque BJM, Trindade F, Herrera H, Chagas da Costa IR, de Oliveira Costa PH, da Costa Fonseca Y, da Silva Valadares RB, Lopes AS. (2024). Metaproteomics revealing microbial diversity and activity in the spontaneous fermentation of maize dough. Food Chem. 435, 137457. 10.1016/j.foodchem.2023.137457

Barbosa Junior D, Gentil K. (2023). Use of Pseudomonas Putida and Pantoea Agllomerans as phosphorus-solubilizing bacteria in latosol. J Interd. Debates, 3 (04), 07–44. 10.51249/jid.v3i04.1065

Cabello P, Luque-Almagro VM, Olaya-Abril A, Sáez LP, Moreno-Vivián C, Roldán MD. (2018). Assimilation of cyanide and cyano-derivatives by Pseudomonas pseudoalcaligenes CECT5344: from omic approaches to biotechnological applications. FEMS Microbiol Lett. 365 (6), 32. 10.1093/femsle/fny032

Dias FMS, Pantoja RK, Gomez JGC, Silva LF. (2023). From degrader to producer: reversing the gallic acid metabolism of Pseudomonas putida KT2440. Int Microbiol. 26, 243–255. 10.1007/s10123-022-00282-5

Ding Z, Fu L, Tie W, Yan Y, Wu C, Hu W, Zhang J. (2019). Extensive Post-Transcriptional Regulation Revealed by Transcriptomic and Proteomic Integrative Analysis in Cassava under Drought. J Agric Food Chem. 67(12), 3521–3534. 10.1021/acs.jafc.9b00014

Dischinger J, Josten M, Szekat C, Sahl H G, Bierbaum G. (2009). Production of the novel two-peptide lantibiotic lichenicidin by Bacillus licheniformis DSM 13. PLoS One. 4, 8, e6788. 10.1371/journal.pone.0006788

Harder C, Hesling E, Botnen S, Lorberau K, Dima B, Bonsdorff-Salminen CV, Niskanen T, Jarvis S, Ouimette A, Hester A, Hobbie E, Taylor A, Kauserud H. (2023). Mycena species can be opportunist-generalist plant root invaders. Environ. Microbiol. 25. 10.1111/1462-2920.16398

Kim JH, Kang SJ, Jung YT, Oh TK, Yoon JH. (2012). Mucilaginibacter lutimaris sp. nov., isolated from a tidal flat sediment. Int J Syst Evol Microbiol. 62, (Pt 3), 515–519. 10.1099/ijs.0.030213-0

Lorenzi AS, Bonatelli ML, Chia MA, Peressim L, Quecine MC. (2022). Opposite Sides of Pantoea agglomerans and Its Associated Commercial Outlook. Microorganisms. 10(10), 2072. 10.3390/microorganisms10102072

Maciel AC, Pena RS, Nascimento LD, Oliveira TA, Chagas-Junior GCA, Lopes AS. (2023) Health exposure risks and bioremediation of cyanide in cassava processing effluents: An overview. J. Water Process. Eng. 55, 2214–7144. 10.1016/j.jwpe.2023.104079

Malmir N, Fard NA, Aminzadeh S, Moghaddassi-Jahromi Z, Mekuto L. (2022). An Overview of Emerging Cyanide Bioremediation Methods. Process. 10(9), 1724. 10.3390/pr10091724

Manabe K, Kageyama Y, Morimoto T, Ozawa T, Sawada K, Endo K, Tohata M, Ara K, Ozaki K, Ogasawara N. (2011). Combined effect of improved cell yield and increased specific productivity enhances recombinant enzyme production in genome-reduced Bacillus subtilis strain MGB874. Appl Environ Microbiol. 77(23), 8370–8381. 10.1128/AEM.06136-11

Medeiros Azevedo T, Aburjaile FF, Ferreira-Neto JRC, Pandolfi V, Benko-Iseppon AM. (2021). The endophytome (plant-associated microbiome): methodological approaches, biological aspects, and biotech applications. World J. Microbiol Biotechnol. 37, 206. 10.1007/s11274-021-03168-2

Mefteh FB, Daoud A, Bouket AC, Alenezi FN, Luptakova L, Rateb ME, Kadri A, Gharsallah N, Belbahri L. (2017). Fungal Root Microbiome from Healthy and Brittle Leaf Diseased Date Palm Trees (Phoenix Dactylifera L.) Reveals a Hidden Untapped Arsenal of Antibacterial and Broad Spectrum Antifungal Secondary Metabolites. Front. Microbiol. 8, 307. 10.3389/fmicb.2017.00307

Mekuto L, Ntwampe SKO, Akcil A. (2016). An integrated biological approach for treatment of cyanidation wastewater. Sci. Total Environ. 571, 711–720. 10.1016/j.scitotenv.2016.07.040

Meyers PR, Rawlings DE, Woods DR, Lindsey GG. (1993). Isolation and characterization of a cyanide dihydratase from Bacillus pumilus C1. J Bacteriol. 175 (19), 6105–6112. https://journals.asm.org/doi/10.1128/jb.175.19.6105-6112.1993

Modesto Junior EN, Chisté RC, Pena RDS. (2019). Oven drying and hot water cooking processes decrease HCN contents of cassava leaves. Food Res Int. 119, 517–523. 10.1016/j.foodres.2019.01.029

Nascimento SV, Costa PH, Herrera H, Caldeira CF, Gastauer M, Ramos SJ, Oliveira G, Valadares RBS. (2022). Proteomic Profiling and Rhizosphere-Associated Microbial Communities Reveal Adaptive Mechanisms of Dioclea apurensis Kunth in Eastern Amazon’s Rehabilitating Minelands. Plants, 11(5), 712. 10.3390/plants11050712

Nwokoro O, Dibua ME. (2014). Degradation of soil cyanide by single and mixed cultures of Pseudomonas stutzeri and Bacillus subtilis. Arh Hig Rada Toksikol. 65, 1, 113–119. 10.2478/10004-1254-65-2014-2449

Okoth RA, Matofari JW, Nduko JM. (2022). Effectiveness of Levilactobacillus brevis fermentation on antinutrients and protein quality of leaves of selected cassava varieties. Appl Food Res. 2, 2, 100134. 10.1016/j.afres.2022.100134

Park JM, Trevor Sewell B, Benedik MJ. (2017). Cyanide bioremediation: the potential of engineered nitrilases. Appl Microbiol Biotechnol, 101, 3029–3042 10.1007/s00253-017-8204-x

Razanamahandry LC, Karoui H, Andrianisa HA, Yacouba H. (2017). Bioremediation of soil and water polluted by cyanide: a review. Afr. J. Environ. Sci. Technol. 11, 272–291. 10.5897/AJEST2016.2264.

Sharma P, Singh SP, Iqbal H.M, Tong YW. (2022). Omics approaches in bioremediation of environmental contaminants: an integrated approach for environmental safety and sustainability. Environ. Res. 211, 113102. 10.1016/j.envres.2022.113102

Sharma M, Akhter Y, Chatterjee S. (2019). A review on remediation of cyanide containing industrial wastes using biological systems with special reference to enzymatic degradation. World J Microbiol Biotechnol. 35, 70. 10.1007/s11274-019-2643-8

Taiwo A, Kudirat O, Tope WA, Abimbola F. (2024). Evaluation of the Potential of Immobilized Cyanide-Degrading Bacteria for the Bioremediation of Cassava Mill Effluent. Jordan J. Biol. Sci. 10.54319/jjbs/170303

Thenmozhi MP, Pracejus B, Novo LAB. (2023). Bioremediation of copper using indigenous fungi Aspergillus species isolated from an abandoned copper mine soil. Chemosphere, 314, 137688. 10.1016/j.chemosphere.2022.137688

Trindade FC, Gastauer M, Ramos SJ, Caldeira CF, Araújo JFD, Oliveira G, Valadares RBS. (2021). Soil metaproteomics as a tool for environmental monitoring of minelands. Forests, 12, 9, 1158. 10.3390/f12091158

Vojcic L, Despotovic D, Martinez R, Maurer K, Schwaneberg U. (2012). An efficient transformation method for Bacillus subtilis DB104. Appl Microbiol Biotechnol. 94, 487–493.10.1007/s00253-012-3987-2

Wang L, Zhang K, Shareen Yin Z, Yu L, Qiu X, Zhou S, Chen R, Wang Q. (2024). Exploring the potential of Aspergillus terreus 2021, WLL-ISO: A dual-functional agent for heavy metal removal and self-aggregation in wastewater treatment. Sep. Purif. Technol. 350, 127964, ISSN 1383-5866, 10.1016/j.seppur.2024.127964.

Wang W, Vignani R, Scali M, Cresti M. (2006). A universal and rapid protocol for protein extraction from recalcitrant plant tissues for proteomic analysis. Electrophoresis, 27, 2782–2786. 10.1002/elps.200500722

Watanabe A, Yano K, Ikebukuro K, Karube I. (1998). Cyanide Hydrolysis in a Cyanide-Degrading Bacterium, Pseudomonas Stutzeri AK61, by Cyanidase. Microbiology. 144, 6, 1677–1682. 10.1099/00221287-144-6-1677

Weimer A, Kohlstedt M, Volke DC, Nikel PI, Wittmann C. (2020). Industrial biotechnology of Pseudomonas putida: advances and prospects. Appl. Microbiol. Biotechnol. 104 (18), 7745–7766. 10.1007/s00253-020-10811-9

Yang L, Fan W, Xu Y. (2020). Metaproteomics insights into traditional fermented foods and beverages. Compr Rev Food Sci Food Saf. 19 (5), 2506–2529. 10.1111/1541-4337.12601

Zango UU, Muhammad II, Sharma V, Sharma AK. (2023). Effective Bioremediation of Cd, Cr, and Pb in Tannery Effluent Using Aspergillus fumigatus and Aspergillus terreus: Synergistic Effects of Using the Two Strains Together. Water Air Soil Pollut. 234, 735. 10.1007/s11270-023-06755-1

Zhou Z, Dong Y, Xia X, Wu S, Huang Y, Liao S, Wang G. (2019). Mucilaginibacter terrenus sp. nov., isolated from manganese mine soil. Int J Syst Evol Microbiol. 69, 10, 3074–3079. 10.1099/ijsem.0.003592

